# Freshwater fish community assessment using eDNA metabarcoding vs. capture-based methods: differences in efficiency and resolution coupled to habitat and ecology

**DOI:** 10.1101/2025.01.08.631834

**Authors:** Manuel Curto, Sofia Batista, Carlos D. Santos, Filipe Ribeiro, Sofia Nogueira, Diogo Ribeiro, Benjamin Prindle, Daniel Licari, Giulia Riccioni, Diogo Dias, Francisco Pina-Martins, Sissel Jentoft, Ana Veríssimo, Maria Judite Alves, Hugo F. Gante

## Abstract

Environmental DNA (eDNA) metabarcoding has revolutionized ecological and environmental research by describing communities without relying on direct observations, making it a powerful, non-invasive, and cost-effective tool in biodiversity monitoring. However, implementation of eDNA as a standard protocol in long-term monitoring programs, that have traditionally relied on capture-based methods, poses challenges in terms of data comparability. Here, we compared freshwater fish communities assessed through eDNA metabarcoding and electrofishing, across 35 sampling sites in the lower Tagus River basin, Portugal. For most species or species-groups analyzed individually (13 out of 17), there was a significant correspondence between electrofishing and eDNA metabarcoding detections. The correspondence was weaker when comparing the number of specimens captured by electrofishing with the number of eDNA metabarcoding reads, with seven out of 13 taxa showing significant relationships. Species richness estimates based on the two methods were very similar at the basin level. The methods yielded significantly different species compositions, although these differences were driven by samples collected in the Tagus main channel, which is wider and has higher flow rates than tributaries. Benthic and shoreline fish communities showed similar species composition in the two methods, but this was not the case for pelagic communities, probably due to the higher water turnover of the pelagic zone and electrofishing inefficiency. Our results highlight the high potential of eDNA metabarcoding as a complementary method to electrofishing for freshwater fish monitoring, though further validation is needed to assess biases related to site-specific hydrological conditions and the ecology of the target species.

## 1. Introduction

In the face of escalating global biodiversity loss and environmental change, the need for more efficient tools for species monitoring is increasingly urgent. The analysis of environmental DNA (eDNA) emerged as a cost-effective and non-invasive method, enabling the identification of target organisms across various life stages using uniform and reproducible criteria (Coble et al., 2019; Evans et al., 2017). Unlike traditional methods relying on direct observation, eDNA analysis does not require taxonomic expertise for morphology-based species identification (Goldberg et al., 2016). eDNA metabarcoding is by far the most frequently used method of eDNA in community analysis, allowing the simultaneous detection of multiple species of different taxonomic groups (Valentini et al., 2016). Despite its high potential, community characterization with eDNA metabarcoding is conditioned by a combination of challenges inherent to DNA barcoding, such as inconsistent taxonomic resolution and or inefficiency of the chosen marker (Valentini et al., 2009; Wilcox et al., 2013), as well as to variability of eDNA release and its persistence in the environment from the timing of its shedding to that of sampling (Pedersen et al., 2015; Yao et al., 2022). While the technical limitations of DNA barcoding are relatively well known, the factors influencing eDNA shedding, persistence and distribution in the environment remain poorly understood (Wang et al., 2021).

Freshwater fish species are characterized by high levels of diversity and provide important ecosystem services (Brummett et al., 2013; Dias et al., 2017). However, they are rapidly being extirpated due to anthropogenic pressures including changes in water flow dynamics, pollution, overfishing, climate change, and the growing presence of invasive species (Revenga et al., 2005; Ricciardi and Rasmussen, 1999; Sala et al., 2000; Suski and Cooke, 2007). Precise quantification of the impacts of these pressures is hindered by difficulties of monitoring fish populations in the wild (Evans and Lamberti, 2018). The established gold standard freshwater fish monitoring techniques are capture-based methods, with electrofishing being the most commonly used (Bonar et al., 2009; Rypel, 2013). However, capture-based methods can be highly selective, and it is often necessary to combine several fishing gears to retrieve an accurate depiction of community diversity and composition (Ruetz III et al., 2007; Schneider, 2000). These methods are also highly demanding in terms of labor, time and equipment, and, in some cases, are still ineffective in capturing rare or elusive species (Bonar et al., 2009; Rypel, 2013). Furthermore, human error and the lack of clear morphological diagnostic traits between closely related taxa make capture-based methods more prone to misidentification (Deiner et al., 2017; Thomsen and Willerslev, 2015). Finally, capture-based methods can be highly invasive, causing harm to animals and the habitats they occupy (Panek and Densmore, 2013), limiting their implementation in monitoring highly threatened species (Deiner et al., 2017; Iknayan et al., 2014; Thomsen and Willerslev, 2015).

eDNA metabarcoding has been proposed as an alternative to capture-based fish monitoring approaches (Evans and Lamberti, 2018). Its efficiency as a monitoring tool has been tested through multiple comparisons with established capture-based approaches, highlighting some of its strengths (Doi et al., 2021; Golpour et al., 2022; Goutte et al., 2020; Jackman et al., 2021; McColl-Gausden et al., 2021; Sard et al., 2019). eDNA metabarcoding is more time- and labor-efficient than capture-based methods (Andres et al., 2023; Evans et al., 2017). For instance, Andres et al. (2023) found that with the same effort, eDNA metabarcoding could screen 25 sites, compared to just nine using seine nets and seven using fyke nets. Furthermore, most eDNA metabarcoding surveys detected similar or even higher species richness compared to capture-based methods (e.g., Andres et al., 2023; Civade et al., 2016; Goutte et al., 2020; Sard et al., 2019; Shaw et al., 2016). Indeed, some studies show that eDNA metabarcoding could retrieve almost all diversity historically recorded at certain locations in a single survey (Antognazza et al., 2021; Hänfling et al., 2016; Nakagawa et al., 2018). This is due to the higher sensitivity of eDNA metabarcoding in detecting rare and elusive species that are typically overlooked by capture-based methods (Gehri et al., 2021). Such feature is particularly important when monitoring taxa of conservation concern, such as endangered and invasive species (Czeglédi et al., 2021).

Nevertheless, some studies showed that eDNA metabarcoding does not produce consistent results across different habitats (Fujii et al., 2019; Li et al., 2022). For example, Civade et al. (2016) found that eDNA metabarcoding outperformed capture-based methods in the tributaries of a lake but not in the lake itself. Similarly, Hallam et al. (2021) described a better performance of eDNA metabarcoding in the freshwater stretches of the Thames River rather than in the estuary. Such inconsistency can result from the heterogeneity of conditions that can impact eDNA shedding, persistence and distribution in the water, including differences in hydrological dynamics (He et al., 2024), water body size (Boivin-Delisle et al., 2021; Moyer et al., 2014), water physicochemical properties (Pilliod et al., 2014; Strickler et al., 2015), and fish ecological and life history traits (Goldberg et al., 2016; Pilliod et al., 2014; Strickler et al., 2015). Variability in eDNA metabarcoding efficiency may be even more pronounced when screening fish diversity at the drainage scale, which may be composed by water bodies of highly contrasting environmental conditions or different fish assemblages with distinct ecological and physiological traits. However, very few studies compared capture-based methods with eDNA metabarcoding at such broad geographical scales.

In this study, we compared the performance of eDNA metabarcoding and electrofishing in assessing of freshwater fish communities in the Tagus basin in Portugal, the longest river system in the Iberian Peninsula. This basin is characterized by high levels of endemism and a high diversity of habitats and hydrological conditions (Collares-Pereira et al., 2021; Sabater et al., 2022). eDNA collection and electrofishing surveys were performed in a large number of sites, covering a variety of environmental and hydrological conditions, and conducted in parallel at each study site to allow direct comparisons of results. In doing so, we assessed the correspondence of fish community metrics between the two methods, and gained insight into the factors contributing to discrepancies, specifically a combination of differences in the environment (channel width) and fish ecology (occupancy of the water column). Through this work, we gained further understanding of the advantages and challenges of eDNA metabarcoding in the context of freshwater fish monitoring at the basin level related to habitat and intrinsic species differences.

## 2. Material and methods

### 2.1 Study area and sampling design

This study was conducted in the lower section of the Tagus River basin, which covers an area of ca. 24,600 km^2^ (Fig. 1). Climate is typically Mediterranean, with hot and dry summers and mild winters (Beck et al., 2018). Average monthly air temperatures range from 7 to 24 °C and annual precipitation is ca. 840 mm (IPMA, 2015). From June 2019 and September 2020, we collected water samples for eDNA analysis and performed electrofishing at 35 sampling sites, covering the Tagus main channel and its tributaries (Fig. 1, Table S1). Seven sampling sites were sampled in two consecutive years (2019 and 2020, Table S1) but these samples were considered independent data points for statistical analysis as the habitat and river flow changed considerably between sampling events.

**Fig. 1.**
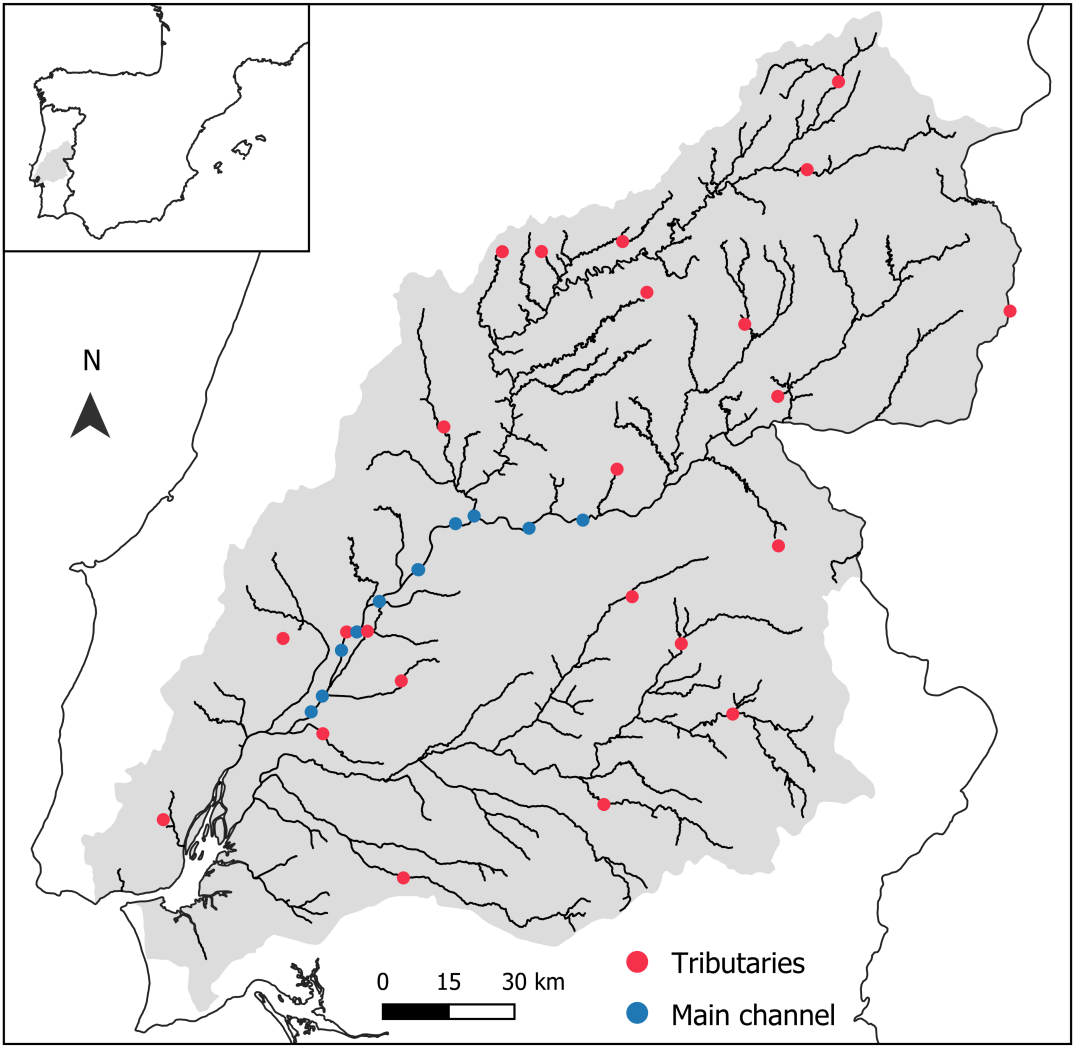
Location of sampling sites in the Tagus River basin. The inset shows the position of the study area within the Iberian Peninsula.

eDNA was obtained by filtering five liters of water from each sampling point. The water was collected with one liter polyethylene plastic bottles along the river channel, ensuring that different habitats were covered (e.g., riffles, pools, areas with aquatic vegetation or plant cover, sandy and/or rocky bottoms). Diligence was applied to avoid suspending bottom sediment or sampling near the water surface, which may release and concentrate suspended organic material, respectively. A boat was used to facilitate water and fish collection from the Tagus main channel. Water samples were filtered on site immediately after collection using nitrocellulose filter capsules (700 cm² and 0.45 µm pore diameter, GoPro™, Company, Location, Country) and a Z-Stream Pump (Merck Millipore, MA, USA) and disposable sterile tubing to minimize possible contamination. After filtering, each capsule was filled with 40 ml of preservation buffer (mixture of 3:1 [v/v] of Lysis Solution to Water Lysis Additive (Sellers et al., 2018) and stored at 4 °C until DNA extraction. To avoid contamination of eDNA samples, we transported water filtering and electrofishing equipment in different vehicles, and sterilized all filtration equipment, including waders, hoses, and gloves, with 10% bleach in-between sampling events. We also produced field negative controls by filtering five liters of bottled commercial water at sampling sites.

Electrofishing was conducted along an 80 m river section at each tributary sampling site using standard gear (EL62 II, Hans Grassl, Germany; discharging 300-600V, 2-5 A, DC). A single anode electrofishing unit was operated by the same person across sampling sites, who waded upstream while ensuring that the different aquatic habitats were surveyed. Sampling sites in the Tagus main channel were surveyed using more powerful electrofishing gear (EL65 II, Hans Grassl, Germany; discharging 350-600V, 8-15 A, DC) following established protocols. After being identified and counted, native species were returned alive to the river, and non-native species were euthanized using an overdose of clove oil (BIOVERT®). All sampling procedures were approved under the research fishing license issued by the national authority for nature conservation (ICNF - Instituto da Conservação da Natureza e das Florestas, I.P.), following national and European regulations on catching and handling wild animals (Decree-Law nr. 92/2019; Directive 2010/63/EU).

### 2.2 DNA Metabarcoding

eDNA extractions were performed using the DNeasy blood and tissue kit (Qiagen, Inc., Hilden, Germany), with the following modifications: Filter capsules were placed at 55 °C for 30 min and vortexed for 1 min to facilitate the release of eDNA from the filter medium. The preservation buffer was extracted from the capsules, transferred to 50 ml falcon tubes, and centrifuged at 5000 rpm for 2 h at room temperature to obtain a pellet. The supernatant was removed, leaving 2 ml of pellet to which 200 µl Proteinase K (10 mg/ml) was added and incubated overnight at 55 °C. After digestion, binding conditions were adjusted by adding 2 ml of AL buffer and 2 ml of absolute ethanol. DNA was eluted in a final volume of 100 μl. The extracted eDNA was quantified using Nanodrop (Thermofisher, Waltham, Massachusetts, USA) and standardized to 10 ng/µl.

DNA metabarcoding was conducted with the MiFish-U primers targeting 170 bp of the 12S mitochondrial gene (Miya et al., 2015) extended with Illumina adapters to facilitate library preparation (Kozich et al., 2013). PCR was performed in a final volume of 20 µl containing 12.5 µl of Qiagen Multiplex PCR Master Mix (Qiagen, Inc., Hilden, Germany), 2 µl of autoclaved H_2_O, 1 µl of each primer (10 µM), and 3 µl of eDNA extract (10 ng/µl). eDNA amplification was performed under the following conditions: 95 °C for 15 min; 35 cycles of denaturation at 98 °C for 20 s, annealing at 60 °C for 15 s, and extension at 72 °C for 15 s; followed by a final extension at 72 °C during 15 min PCR negative controls were added by using water as template. Eight replicates were performed per sample to reduce amplification stochasticity associated with PCR. Amplification success was assessed by 2% agarose gel electrophoresis and products were quantified using Nanodrop. PCR replicates were then pooled (2.5 µl per sample) and the resulting solution used as template for indexing. This was done through a 10-cycle PCR reaction with KAPA HiFi HotStart Kit (Roche Molecular Systems, USA) using the following temperature profile: 95 °C for 3 min, 10 cycles of denaturation at 95 °C for 30 s, annealing at 55 °C for 30 s, and extension at 72 °C for 30 s, followed by final extension step at 72 °C for 5 min. Indexed PCR product size was measured by agarose gel electrophoresis and cleaned with Ampure beads (Beckman Coulter, Brea, California, USA). The purified amplicons were quantified using Nanodrop, normalized at 15 nM and pooled equimolarly. The final pool was checked for size and quantity on a TapeStation (Aligent, USA) and validated by quantitative PCR assay with primers specific to the Illumina adapters. The final pool was sequenced using an Illumina MiSeq (Illumina, San Diego, CA, USA) and the v2 kit (250 bp paired-end reads). Each sample was sequenced with a target depth of 10k reads. Indexing and sequencing was conducted as a service at the Centre for Molecular Analysis (CTM) from CIBIO/BIOPOLIS.

### 2.3 Sequence data analysis

Illumina adapters and amplification primers were removed using Cutadapt v.3.4 (Martin, 2011). Only sequence reads that started with the forward primer, ended with the reverse primer, and had a length greater than 80 bp were analyzed. Sequence data were filtered to produce quality reads through the following steps: truncating reads to 165 bp, merging paired-end reads, removing chimeras, and inferring amplicon sequence variants (ASVs) using run-specific error rates. These procedures were performed using R package DADA2 v.1.20 (Callahan et al., 2016). ASV counts per sample were stored in a matrix. These were then corrected and filtered to minimize false positives related to tag-jumping and contamination. A correction factor of 5% was applied using the equation from Hambäck et al. (2021) to reduce the impact of tag-jumping signals. This value was obtained empirically by evaluating the number of reads originating from species targeted in other projects that shared the same Illumina run and used the same marker. These projects targeted marine bony and cartilaginous fish species that are not found in the Tagus basin. Signals possibly originating from contamination were controlled by subtracting the maximum read count per taxon found in the negative controls.

Taxonomic assignment of the ASVs was performed using the megaBLAST module from BLAST+ 2.12.0 (McGinnis and Madden, 2004) on the databases MitoFish v3.85 and GenBank Release 246.0 (Sayers et al., 2021). BLAST outputs were filtered according to e-value, identity (ID) and query coverage following Curto (2022). Only ASVs with e-value <1e^-10^, query coverage and ID >90% were kept. Full lineage information from each match was obtained using TaxonKit v0.9.0 (Shen and Ren, 2021). In case of multiple matches with the same e-value, ID and query coverage, the most recent common taxon was chosen. Assignments at genus or family levels were further evaluated through phylogenetic analyses using publicly available sequences of the 12S gene. These sequences were downloaded from GenBank and aligned with the corresponding ASVs using muscle (Edgar, 2004) as implemented in Aliview v1.28 (Larsson, 2014). The optimal nucleotide substitution model was obtained using jModelTest 2.1.10 (Posada, 2008) implementing the Akaike information criterion. Phylogenetic trees were built using the software RAxML v8.2.12 (Stamatakis, 2014) using 1000 bootstrap replicates to calculate node support. The output was visualized using FigTree v1.4.4 (Rambaut and Drummond, 2010) and, in case the ASV fell within a monophyletic clade, it was assigned to the taxon of the remaining members of the clade.

### 2.4 Statistical analysis

The results of fish community assessment through electrofishing and eDNA metabarcoding were compared with three main analytical approaches: (1) Species-specific Generalized Linear Models (GLM); (2) Non-metric Multidimensional Scaling (NMDS) of fish assemblages; (3) Rarefaction curves of species richness.

GLMs were used to evaluate the similarity of results from both methods for species and species-groups treated separately. For this analysis, we excluded taxa that appeared in fewer than 10% of the samples, as their data were insufficient for modeling purposes. We first fitted binomial GLMs, with electrofishing detection (detected *vs* not detected) as dependent variable and eDNA metabarcoding detection (detected *vs* not detected) as predictor. These models were fitted using Firth’s penalized likelihood estimation using the function logistf of the logistf R package v1.26.0 (Ploner, 2010), as detections for several taxa matched almost completely between electrofishing and eDNA metabarcoding. For these taxa showing a significant relationship between electrofishing and eDNA detections, we further investigated possible relationships between the number of specimens captured and the number of reads in eDNA metabarcoding. For this purpose, we fitted GLMs with a negative binomial distribution using the function glm.nb of the MASS R v7.3-60 package (Venables, 2002). The parameter theta of negative binomial distributions was adjusted automatically by the glm.nb function to cope with high number of zeros and low counts of electrofishing data.

The NMDS analysis had the purpose of providing a general perspective of the existing differences in fish assemblages inferred from electrofishing and eDNA metabarcoding in the array of sampling points. This analysis was conducted with functions from the Vegan R package v2.6-4 (Oksanen et al., 2013). First, we standardized electrofishing and eDNA data with the decostand function, then we built a Bray-Curtis dissimilarity matrix with the vegdist function, and finally we performed the NMDS with the metaMDS function. We additionally tested the overall differences between methods (electrofishing and eDNA) using an Analysis of Similarities (ANOSIM), for which we used the function anosim of Vegan package. We repeated NMDS and ANOSIM analyses for subsets of data from sampling locations in the Tagus main channel and in tributaries, and for groups of species categorized by their ecological habitat preferences: pelagic, benthic and shoreline (classifications followed Collares-Pereira et al., 2021). These analyses aimed to examine the contribution of river configuration (e.g., width and depth) and species ecology to the dissimilarity in results of electrofishing and eDNA metabarcoding.

Finally, the rarefaction curves of species richness had the purpose of comparing the efficiency of electrofishing and eDNA metabarcoding in detecting additional species as the number of sampling points (i.e., sampling effort) increased. With this approach, we were able to compare species richness estimates provided by each method and the sampling effort necessary to reach those estimates. Besides the general comparison of sampling methods, we built specific curves for sampling locations of the Tagus main channel and those of its tributaries, to test if river size could interfere with the performance of the sampling method. Rarefaction curves were built in R with the specaccum function of the Vegan package with 10,000 permutations. Species richness rarefaction curves were complemented with regression analyses to examine the relationship between species richness estimated by electrofishing and that of eDNA metabarcoding. We first tested for a significant relationship between these two variables by fitting a GLM with a Poisson distribution and log link function, where eDNA species richness was used as a predictor of electrofishing species richness. Then, to test if the two variables were directly proportional, we fitted a null model for electrofishing species richness with log-transformed eDNA species richness as an offset term. This GLM also employed a Poisson distribution with a log link function. In this model, a β coefficient not significantly different from zero would indicate that electrofishing species richness and eDNA species richness are directly proportional. Both GLMs were fitted using the glm function from the stats package in R v4.3-3 (R Core Team, 2023).

## 3. Results

A total of 12,022 individuals, representing 31 taxa in 19 fish families, were collected in the 42 electrofishing surveys (Table S2). The eastern mosquitofish (*Gambusia holbrooki*), a non-native species, was by far the most abundant fish collected (3,755 individuals). Other fishes that surpassed 1,000 collected individuals were all native taxa: the big-scale sand smelt (*Atherina boyeri*), the southern Iberian spined loach (*Cobitis paludica*) and the species-group *Squalius* spp., composed by individuals of the *S. alburnoides* complex and the Southern Iberian chub (*S. pyrenaicus*) (Table S2). Regarding the frequency of occurrence over the different sampling sites, the barbels (*Luciobarbus* spp., composed by *L. bocagei*, *L. comizo* and *L. steindachneri*) and the eastern mosquitofish (*G. holbrooki*) were the most frequently collected (Table S2).

For eDNA metabarcoding, 1,675,638 raw reads were produced (average of 18,413 per sample), of which 1,460,683 passed the adapter and primer trimming step (Table S1). DADA2 kept 1,244,640 reads grouped into 1,680 ASVs, with a mean of 33 per sampling point (‘station’ in Table S1). 1,604 ASVs (1,242,648 reads) were assigned with a minimum of 90% identity match to 204 taxa (Table S1). Most ASVs were assigned to Bacteria (1,117), however this only corresponded to 3.8% of the total number of reads. Conversely, 156 ASVs were assigned to freshwater fish species, representing 91,9% of the total reads. The remaining reads were mostly assigned to other vertebrate groups.

After applying all filters and corrections based on negative controls and tag-jumping, eDNA metabarcoding identified 71 vertebrate taxa, of which 31 freshwater and migratory fishes, 14 mammals (including humans), 14 birds, seven amphibians and one testudine (Table S3). For freshwater and migratory fishes, phylogenetic analyses determined the assignment of three fish taxa: European eel (*Anguilla Anguilla*), the Eurasian carp (*Cyprinus carpio*), and flathead grey mullet (*Mugil cephalus*). For *Squalius* the tree lacked resolution to delineate many of the species and therefore was not informative to clarify taxonomic ambiguities. However, all ASVs classified at the genus level showed equal best matches to *S. pyrenaicus* and *S. alburnoides* and the final classification considered both species (*Squalius pyrenaicus/alburnoides).* In eight other taxa, ASVs formed monophyletic groups with two species (Fig. S1). Six of these pairs exist in the Tagus basin (*Chelon labrosus/ramada*, *Luciobarbus bocagei/comizo*, *Alosa alosa/fallax, Carassius auratus/gibelio,* and *Lampetra planeri/auremensis*), thus the taxonomic assignment considered both species (Table S2). *Sander* and *Salmo* ASVs were assigned to Eurasian pikeperch (*Sander lucioperca*) and brown trout (*Salmo trutta*), respectively, as the alternative pairs (estuarine perch, *Sander marinus*, and Black Sea salmon, *Salmo labrax*) are not present in Portuguese waters and rarely enter rivers (Freyhof and Kottelat, 2007; Gilbert, 1993).

From the 31 freshwater fish taxa identified, the species-group *Squalius* spp. had the highest eDNA read count, followed by the mullets (*Chelon* spp.), the Pyrenean gudgeon (*Gobio lozanoi*) and the barbels (*Luciobarbus* spp.) (Table S2). Similar to results from electrofishing, the barbels and the eastern mosquitofish showed the highest frequency of occurrence across the sampling sites (Table S2).

In general, there was reasonable correspondence between the results of electrofishing and eDNA metabarcoding, both in terms of frequency of occurrence, and in the abundance of captures and eDNA read counts (Fig. 2, Table S2). The ratio “number of eDNA reads: number fish captured” ranged from ca. 50 to 300 in the vast majority of cases (Table S2). Compared to this norm, some taxa are clear outliers indicating under- or overperformance of one of the methods. eDNA metabarcoding comparatively underperformed for the big-scale sand smelt and the eastern mosquitofish, and overperformed for all mullets (flathead grey mullet, *Mugil cephalus*, and species-group *Chelon* spp., which includes *C. labrosus* and *C. ramada*) and for the Eurasian carp (*Cyprinus carpio*) (Fig. 2, Table S2). A similar comparison based on frequency of occurrence supports a clear overperformance of eDNA metabarcoding in the detection of the flathead grey mullet. For these comparisons, we excluded taxa with frequency of occurrence below 0.1% and one outlier sampling point for the southern Iberian spined loach, where we caught 80% of the individuals of this species. Abundance mismatches between electrofishing and eDNA metabarcoding across the sampling points were evident for several species and species-groups, such as the barbels group (*Luciobarbus* spp.), the eastern mosquitofish, the pumpkinseed sunfish (*Lepomis gibbosus*) and the *Squalius* spp. group (Fig. 2).

**Fig. 2.**
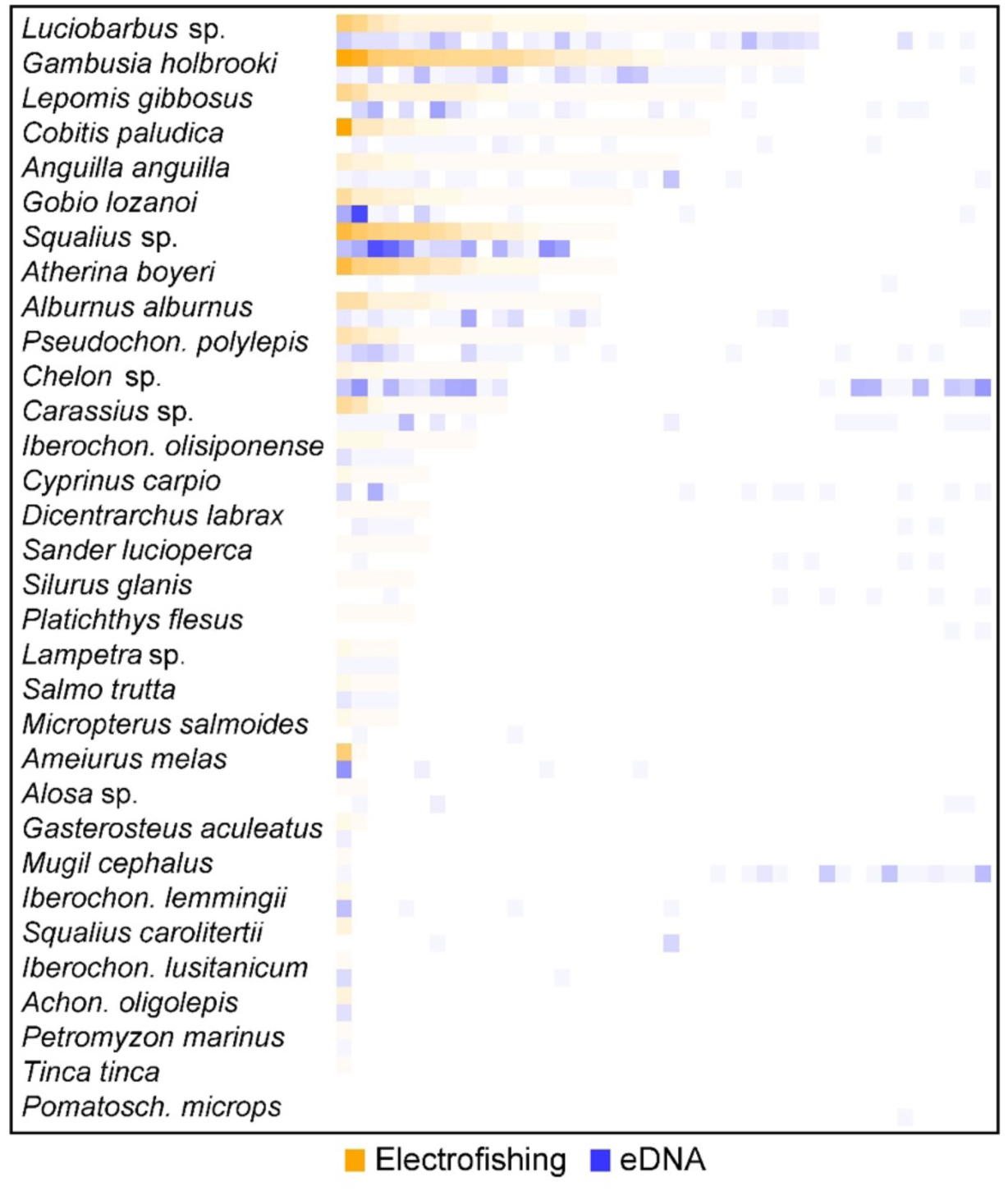
Heatmap contrasting the number of specimens captured by electrofishing with the number of reads from eDNA metabarcoding in each sampling site. Color scale is logarithmic to minimize the influence of extreme observations. Species are listed in descending order of frequency of occurrence in electrofishing samples. Sampling sites are ordered horizontally per species from the highest to lowest number of captures in electrofishing. Abbreviations: *Pseudochon*. – *Pseudochondrostoma*; *Iberochon*. – *Iberochondrostoma*; *Achon*. – *Achondrostoma*; *Pomatosch*. – *Pomatoschistus*

The common goby (*Pomatoschistus microps)* was detected in eDNA metabarcoding but not collected with electrofishing ((Fig. 2, Table S1). Conversely, the Eurasian tench (*Tinca tinca*) was captured in one sampling site but not detected in eDNA metabarcoding (Fig. 2, Table S2).

Consistency between sampling methods (electrofishing and eDNA metabarcoding) was evident from the results of the regression models (Table 1). For the vast majority of species and species-groups tested (13 out of 17), we found a significant relationship between electrofishing and eDNA detections (Table 1). Seven of these 13 cases also showed a significant relationship between the number of fish collected and the number of eDNA reads per species and per sampling point (Table 2).

**Table 1.**
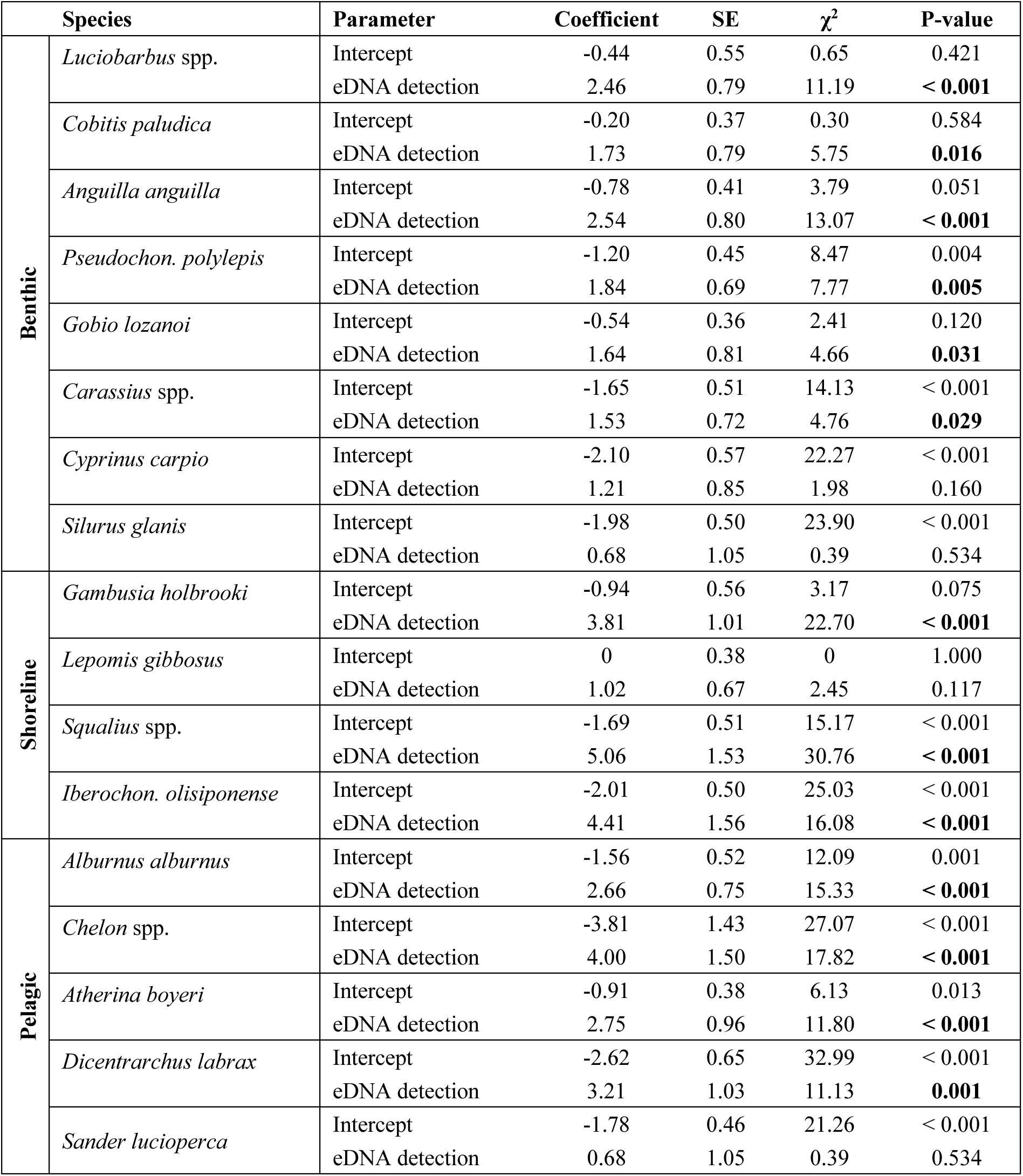
Summary statistics of Binomial Generalized Linear Models (GLM) testing the relationship between fish species detection in electrofishing and eDNA metabarcoding. SE - Standard error; χ^2^ – Qui-square statistics. Significant relationships are shown in bold. Species occurring in less than 10% of the electrofishing and eDNA samples were excluded. Within each species group, the species are listed in descending order of frequency of occurrence in samples. Abbreviations: *Pseudochon*. – *Pseudochondrostoma*; *Iberochon*. – *Iberochondrostoma*.

**Table 2.** Summary statistics of Generalized Linear Models (GLM) testing the relationship between the number of fish captured with electrofishing and the number of detections in eDNA metabarcoding. Models were fitted with a negative binomial distribution. SE - Standard error; Z – Z statistics. Significant relationships are shown in bold. Species for without significant relationships in Binomial GLMs (table 1) or occurring in less than 10% of the electrofishing and eDNA samples are not represented. Within each species group, the species are listed in descending order of frequency of occurrence in samples. Abbreviations: *Pseudochon*. – *Pseudochondrostoma*; *Iberochon*. – *Iberochondrostoma*.

| | Species | Parameter | Coefficient | SE | $\chi^2$ | P-value |
| --- | --- | --- | --- | --- | --- | --- |
| Benthic | *Luciobarbus* spp. | Intercept | 2.08 | 0.31 | 6.60 | < 0.001 |
|  |  | eDNA read count | 3.2x10<sup>4</sup> | 8.9x10<sup>5</sup> | 3.58 | < **0.001** |
|  | *Cobitis paludica* | Intercept | 3.61 | 0.45 | 7.94 | < 0.001 |
|  |  | eDNA read count | -3.3x10<sup>4</sup> | 9.2x10<sup>4</sup> | -0.37 | 0.714 |
|  | *Anguilla anguilla* | Intercept | 1.53 | 0.32 | 4.75 | < 0.001 |
|  |  | eDNA read count | 3.7x10<sup>4</sup> | 2.3x10<sup>4</sup> | 1.65 | 0.098 |
|  | *Pseudochon. polylepis* | Intercept | -0.03 | 0.37 | -0.08 | 0.934 |
|  |  | eDNA read count | 9.6x10<sup>4</sup> | 1.8x10<sup>4</sup> | 5.48 | < **0.001** |
|  | *Gobio lozanoi* | Intercept | 1.70 | 0.40 | 4.25 | < 0.001 |
|  |  | eDNA read count | 1x10<sup>4</sup> | 3.1x10<sup>5</sup> | 3.06 | **0.002** |
|  | *Carassius* spp. | Intercept | 0.85 | 0.61 | 1.40 | 0.162 |
|  |  | eDNA read count | 1.6x10<sup>3</sup> | 3.5x10<sup>4</sup> | 4.46 | < **0.001** |
| Shoreline | *Gambusia holbrooki* | Intercept | 4.21 | 0.36 | 11.54 | < 0.001 |
|  |  | eDNA read count | 1.5x10<sup>4</sup> | 1.1x10<sup>4</sup> | 1.31 | 0.189 |
|  | *Squalius* spp. | Intercept | 2.37 | 0.50 | 4.70 | < 0.001 |
|  |  | eDNA read count | 1.4x10<sup>4</sup> | 3.3x10<sup>5</sup> | 4.32 | < **0.001** |
|  | *Iberochon. olisiponense* | Intercept | -0.94 | 0.49 | -1.91 | 0.057 |
|  |  | eDNA read count | 5.5x10<sup>3</sup> | 7.9x10<sup>4</sup> | 6.93 | < **0.001** |
| Pelagic | *Alburnus alburnus* | Intercept | 1.90 | 0.46 | 4.12 | < 0.001 |
|  |  | eDNA read count | 3x10<sup>4</sup> | 1.7x10<sup>4</sup> | 1.80 | 0.073 |
|  | *Chelon* spp. | Intercept | -0.19 | 0.54 | -0.36 | 0.720 |
|  |  | eDNA read count | 1.6x10<sup>4</sup> | 7x10<sup>5</sup> | 2.34 | **0.019** |
|  | *Atherina boyeri* | Intercept | 3.45 | 0.53 | 6.55 | < 0.001 |
|  |  | eDNA read count | 2.0x10<sup>4</sup> | 1.2x10<sup>3</sup> | 0.16 | 0.872 |
|  | *Dicentrarchus labrax* | Intercept | -1.40 | 0.53 | -2.65 | 0.008 |
|  |  | eDNA read count | 2.2x10<sup>3</sup> | 1.1x10<sup>3</sup> | 1.94 | 0.053 |

NMDS analysis revealed significant overall dissimilarities in the species composition of the sampling points for the two methods (electrofishing and eDNA metabarcoding) (Fig. 3). However, when splitting the dataset between samples collected in the Tagus main channel and in its tributaries, it became evident that the dissimilarities between methods were mostly generated by the main channel subset of sampling points (Fig. 3), while distances between fish assemblages sampled with electrofishing and eDNA metabarcoding were relatively low in the tributaries (Fig. 3). When analyzing fish species grouped according to their habitat preferences, we found that while benthic and the shoreline communities were very similar when surveyed with electrofishing or eDNA metabarcoding, pelagic communities sampled with the two methods were significantly different (Fig. 3).

**Fig. 3.**
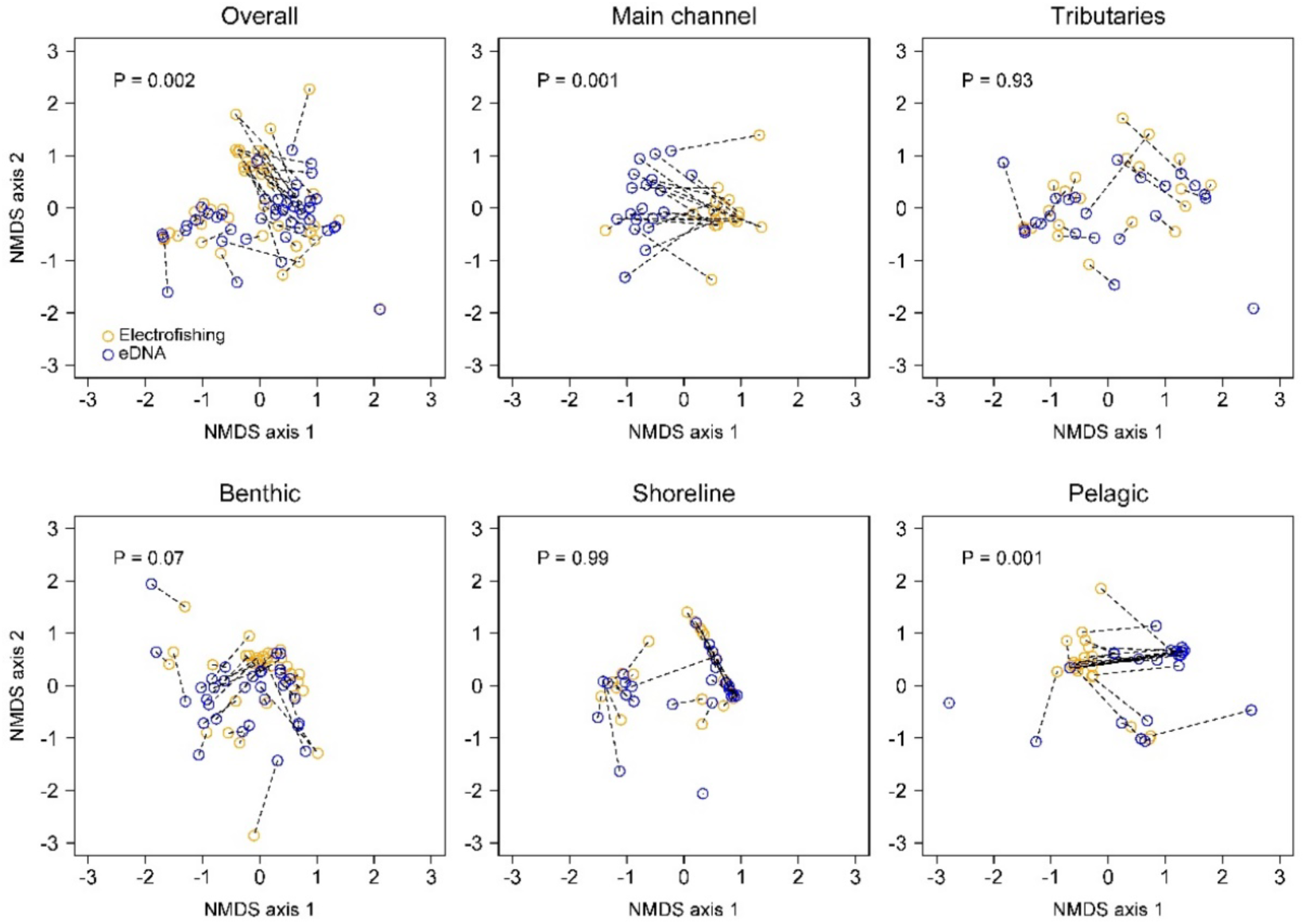
Non-metric Multidimensional Scaling (NMDS) biplots contrasting fish assemblages sampled with electrofishing and eDNA metabarcoding for each sampling site. The results of the two methods for each sampling site are linked by dashed lines. P-values shown inside the plots indicate statistical significance of differences between sampling methods (electrofishing and eDNA metabarcoding) based on an Analysis of Similarities (ANOSIM). Bottom panels show results for species-groups with distinct habitat preferences. Benthic: *Anguilla anguilla*, *Ameiurus melas*, *Carassius* spp., *Cobitis paludica*, *Cyprinus carpio*, *Gobio lozanoi*, *Iberochondrostoma lemmingii*, *Iberochondrostoma lusitanicum*, *Lampetra* spp., *Luciobarbus* spp., *Petromyzon marinus*, *Platichthys flesus*, *Pseudochondrostoma polylepis*, *Silurus glanis*; Shoreline: *Achondrostoma oligolepis*, *Gambusia holbrooki*, *Gasterosteus aculeatus*, *Iberochondrostoma olisiponense*, *Lepomis gibbosus*, *Squalius carolitertii*, *Squalius* spp.; Pelagic: *Alburnus alburnus*, *Alosa* spp., *Atherina boyeri*, *Chelon* spp., *Dicentrarchus labrax*, *Micropterus salmoides*, *Mugil cephalus*, *Salmo trutta*, *Sander lucioperca*.

Despite the differences between electrofishing and eDNA metabarcoding results at the species and assemblage levels, these two methods produced similar inferences of species richness. Specifically, the rarefaction curves for the two methods converged to similar species richness estimates, with overlapping confidence intervals, and showed an almost identical trend demonstrating similar power to discover new species as the sampling effort increases (Fig. 4). Rarefaction curves for the two methods overlapped less when the dataset was split into main channel and tributary subsets, although the differences between the curves were not significant in either data subset (Fig. 4). These results were supported by regression analysis results showing highly significant relationship between electrofishing and eDNA metabarcoding species richness (Poisson GLM: β = 0.054, SE = 0.015, p < 0.001). Additionally, the null model, where log-transformed eDNA species richness was used as an offset for electrofishing species richness, indicated that these two variables were directly proportional (β = 0.079, SE = 0.057, p = 0.17; see Methods for statistical details). This suggests that increases in species richness detected by electrofishing were closely mirrored by those detected by eDNA metabarcoding.

**Fig. 4.**
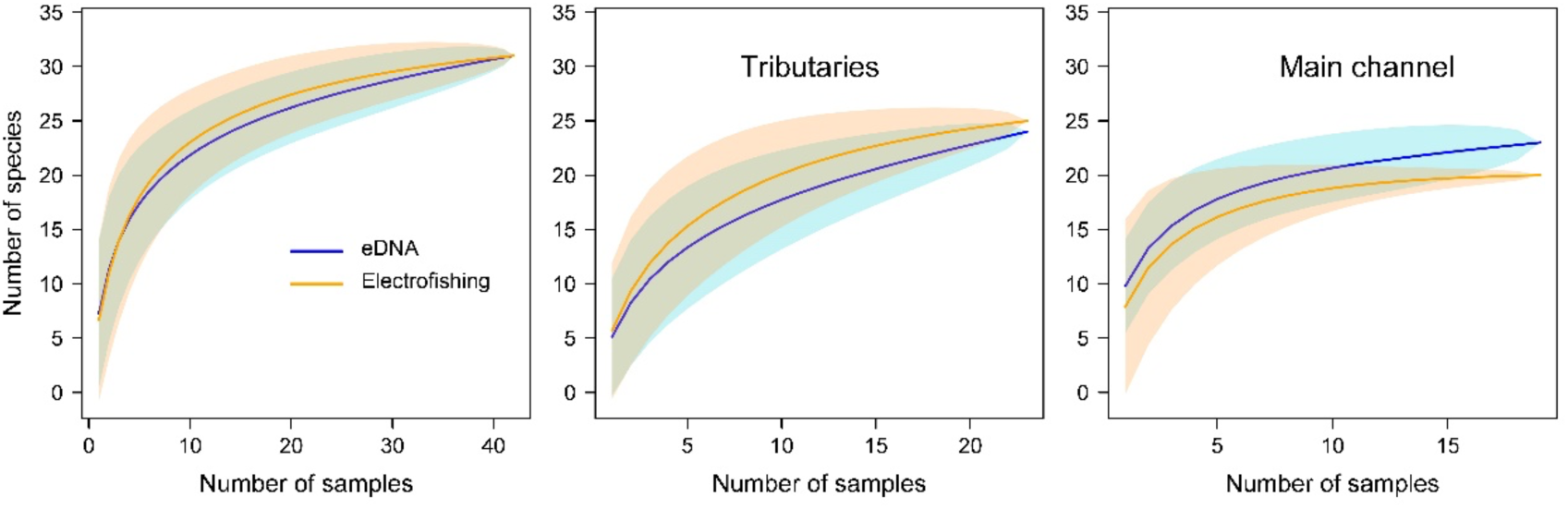
Rarefaction curves of species richness resulting from electrofishing and eDNA metabarcoding detections. Shading represents 95% confidence intervals.

## 4. Discussion

The simple, fast, non-invasive, and resource-efficient fieldwork protocol associated with water collection for eDNA metabarcoding, coupled to its high sensitivity in species detection and streamlined species identification, makes it a particularly attractive method for monitoring freshwater fish diversity at large geographical scales (Andres et al., 2023; Nakagawa et al., 2018), as it is the case here for the lower Tagus Basin in Portugal. Furthermore, the presence of specialists (e.g., taxonomists and bioinformaticians) is not required during eDNA collection but rather for sample processing, data analysis, and interpretation, which can be performed later, facilitating the management of human resources compared to capture-based methods. We found an overall correspondence between eDNA metabarcoding and electrofishing, both in terms of species richness and species detection ability, indeed suggesting that eDNA metabarcoding is a viable and comparable option for monitoring fish communities. Moreover, the heterogeneity of conditions found within the studied basin, allowed us to identify the cases where between-method correspondence falls short, especially in terms of community composition, highlighting some of the factors affecting species detection ability in eDNA metabarcoding.

Detection efficiency was congruent between methods for most taxa. Among the 17 taxa that were abundant enough to be analyzed individually, 13 showed significant concordance between eDNA and electrofishing detections across the sampling points. However, the correspondence of abundance proxies between electrofishing and eDNA metabarcoding was not as high. Indeed, seven taxa (out of the 13 with consistent detection between methods) showed a positive and significant relationship between electrofishing-based fish abundance and eDNA read counts. Notably, eDNA read counts were particularly low compared to the number of specimens caught for the big-scale sand smelt and the eastern mosquitofish, suggesting eDNA metabarcoding underperformance, while the opposite was observed for the mullets and the common carp, which tend to be well-documented in eDNA studies (Pont et al., 2018; Van Driessche et al., 2023a). These detection discrepancies between species may reflect simple differences in body size or any other physiological traits that impact eDNA shedding and decay rates, and consequently detection probabilities.

Even when correlations between eDNA read counts and number of captured individuals are found,, these tend to be weak (Boivin-Delisle et al., 2021; Di Muri et al., 2020; Hallam et al., 2021; Kelly et al., 2019). Such deviations between read count and number of captured individuals can be explained by multiple factors including gear selectivity (Evans and Lamberti, 2018; Gehri et al., 2021), differential efficiency of PCR amplification across taxonomic groups (Goldberg et al., 2016; Hallam et al., 2021), and the hydrological, physicochemical, and physiological and ecological factors that influence the release, distribution and decay of eDNA in the water (Andruszkiewicz Allan et al., 2021; Deiner et al., 2017; Maruyama et al., 2014; Pilliod et al., 2014; Strickler et al., 2015; Thalinger et al., 2021). The way some of these factors impact eDNA metabarcoding read counts is still unclear, requiring additional benchmarking for robust and reliable proxy for fish abundance or biomass. Nonetheless, analytical corrections such as allometric scaling are promising (Yates et al., 2021).

In this study, eDNA metabarcoding and electrofishing recovered similar levels of species richness at the basin level. Such a result was obtained despite a relatively high sampling effort being put into place for electrofishing in the main river channel, increasing its ability to recover a relatively high proportion of fish diversity. This included the sampling of different habitats within each sampling site and the adjustment of sampling effort according to channel width (Bernardo et al., 2008). Conversely, this adjustment was not made for eDNA metabarcoding, as we filtered the same volume of water and ran the same number of PCR replicates in all sites independently of river type. In stretches of the basin with greater channel width and depth, eDNA sensitivity is lower due to increased dilution, requiring the sampling of larger volumes of water or increased PCR replication (Boivin-Delisle et al., 2021; Curtis et al., 2021; Moyer et al., 2014; Pont et al., 2018). Had we adjusted the eDNA sampling effort, it is likely we would have detected additional biodiversity. Nonetheless, these results show that even a standardized protocol applied across different sampling sites is sufficient for an initial assessment of highly heterogeneous systems. Future efforts can then refine the eDNA field and lab protocols based on the specific characteristics of each sampling site.

We found significant differences in local community composition delivered from both methods. Interestingly, by sub-setting the data according to channel width, we verified that these differences only remain when considering survey locations in the main channel. Similar results have been reported in other studies comparing water bodies with contrasting volumes (Civade et al., 2016; Hallam et al., 2021). Such differences may be explained by multiple factors that can impact detection sensitivity with both methods, such as habitat complexity, water flow patterns across the channel, and insufficient sampling effort. Indeed, some portions of the Tagus main channel are highly variable in terms of habitats and hydrological conditions. In such river sections, both individual fish and their eDNA are likely distributed more heterogeneously (Hallam et al., 2021). While our results support the ability of eDNA metabarcoding in providing accurate snapshots of fish communities in smaller rivers (Van Driessche et al., 2023b), eDNA may remain localized in river backwaters and side pools, be captured by sediment or river substrate (Jerde et al., 2016), or become unevenly distributed due to currents (He et al., 2024; Lawson Handley et al., 2019) and thermal stratification (Jeunen et al., 2023; Littlefair et al., 2023) in the artificial reservoirs located in the main channel, which are akin to lentic environments. Likewise, some species may associate with discrete conditions in such large areas, such as patches of aquatic vegetation, or areas of lower/higher water currents (e.g., *Iberochondrostoma olisiponense* (Veríssimo et al., 2018)). Such heterogeneity can exacerbate the differences in community composition retrieved by eDNA metabarcoding and capture-based methods if the waterbody is not thoroughly sampled (He et al., 2024). Although an effort was made to sample different habitats with both electrofishing and eDNA metabarcoding, we may have not covered all of them efficiently or similarly, thereby biasing the portion of the fish community sampled by each method. In the tributaries, such high heterogeneity in hydrological and habitat complexity is less likely to occur given the smaller spatial scales. Consistent with higher sensitivity in smaller rivers, eDNA metabarcoding detected an additional species in tributaries that were not collected by electrofishing.

Fish ecotype also contributed to community composition differences between methods for species occupying the water column. Pelagic species showed significant differences between eDNA metabarcoding and electrofishing, but benthic and shoreline species did not. This may be explained by the difficulty of capturing some of the fast-swimming species living in the pelagic zone with electrofishing (Mahon, 1980). In fact, the absence of European seabass (*Dicentrarchus labrax)* in our results demonstrates this, since this fish is a fast swimmer and may easily escape at the slightest disturbance of the water, being common in the main steam of the Tagus (Almeida et al., 2024). These results may also be related to the way eDNA from different groups of organisms travels within the system. Benthic and littoral organisms are in closer contact with the substrate and therefore their eDNA may easily become trapped in the bottom, or travel shorter distances (Jerde et al., 2016). Conversely, eDNA from pelagic organisms is more likely to remain longer periods in the water column traveling longer distances, especially in river sections with fast current and high discharge rates (Deiner and Altermatt, 2014; Van Driessche et al., 2023a). Thus, some of the detections of pelagic fish species may stem from animals living upstream of the sampling sites, which could not have been detected with electrofishing.

Two taxa were only detected by one of the sampling methods. The common goby was detected exclusively by eDNA metabarcoding, while the Eurasian tench was collected only by electrofishing. In fact, the Eurasian tench that was found to shed comparatively less eDNA, which might decrease detection sensitivity (Caza-Allard et al., 2022). These are relatively rare taxa making their detectability challenging for either method (Fujii et al., 2019).

## 5. Conclusion

The overall similarity in performance between both methods reinforces the value of eDNA metabarcoding as a monitoring tool for freshwater fish species at the basin level. However, this study also highlights that eDNA detection sensitivity may be variable, as it depends on environmental factors and fish ecology. Some of the conditions where eDNA metabarcoding underperformed compared to electrofishing can be addressed. For example, portions of the basin with wider river channels can be more thoroughly sampled. Biases related with species ecology and eDNA transport and decay are more challenging, yet can be handled with proper validation and potentially corrected with predictive models based on prior data (Carraro et al., 2023). Thus, before any implementation of eDNA sampling as the standard protocol (e.g., in cases where species diversity is a key focus), a transition phase combining traditional capture-based approaches and eDNA metabarcoding is warranted. Fundamental research on this topic is crucial to pave the way for eDNA approaches in biodiversity and conservation, particularly regarding the environmental and biological variables that determine species and habitat differences in eDNA shedding, distribution, and decay.

## Acknowledgments

Electrofishing was conducted under the fishing credentials 268-272/2019/CAPT issued by ICNF. Furthermore, we thank Francisco Pinto and João Lobo for providing their boats while fishing in some of the localities of the Tagus main channel. Library preparation and sequencing was done by the molecular analysis services at CIBIO-BIOPOLIS. The computational analyses were performed on the Saga Cluster owned by the UiO and the Norwegian metacenter for High Performance Computing (NOTUR) and operated by the UiO Department for Research Computing (https://www.hpc.uio.no).

## CRediT authorship contribution statement

**Manuel Curto:** Conceptualization, Data curation, Formal analysis, Investigation, Methodology, Supervision, Validation, Visualization, Writing – original draft. **Sofia Batista:** Conceptualization, Data curation, Formal analysis, Investigation, Methodology, Validation, Visualization, Writing – original draft. **Carlos D. Santos**: Conceptualization, Data curation, Formal analysis, Investigation, Methodology, Supervision, Validation, Visualization, Funding acquisition, Project administration, Writing – original draft. **Filipe Ribeiro:** Data curation, Funding acquisition, Investigation, Methodology, Project administration, Resources, Supervision, Validation, Visualization, Writing – review and editing. **Sofia Nogueira:** Data curation, Methodology, Validation, Visualization, Writing – review and editing. **Diogo Ribeiro:** Data curation, Methodology, Validation, Visualization, Writing – review and editing. **Benjamin Prindle**: Methodology, Validation, Visualization, Writing – review and editing. **Daniel Licari:** Methodology, Validation, Visualization, Writing – review and editing. **Giulia Riccioni:** Investigation, Methodology, Validation, Visualization, Writing – review and editing. **Diogo Dias:** Methodology, Validation, Visualization, Writing – review and editing. **Francisco Pina-Martins:** Data curation, Supervision, Validation, Visualization, Writing – review and editing. **Sissel Jentoft:** Funding acquisition, Resources, Validation, Visualization, Writing – review and editing. **Ana Veríssimo:** Data curation, Funding acquisition, Investigation, Methodology, Project administration, Resources, Supervision, Validation, Visualization, Writing – review and editing. **Maria Judite Alves:** Funding acquisition, Investigation, Methodology, Project administration, Resources, Supervision, Validation, Visualization, Writing – review and editing. **Hugo F. Gante**: Conceptualization, Formal analysis, Funding acquisition, Investigation, Methodology, Project administration, Resources, Supervision, Validation, Visualization, Writing – review and editing

## Data availability

Raw sequence data is deposited in the SRA database from NCBI under the project PRJNA1171386. Further data will be made available on request.

