## Supplementary for "Freshwater fish community assessment using eDNA metabarcoding vs. capture-based methods: differences in efficiency and resolution coupled to habitat and ecology"

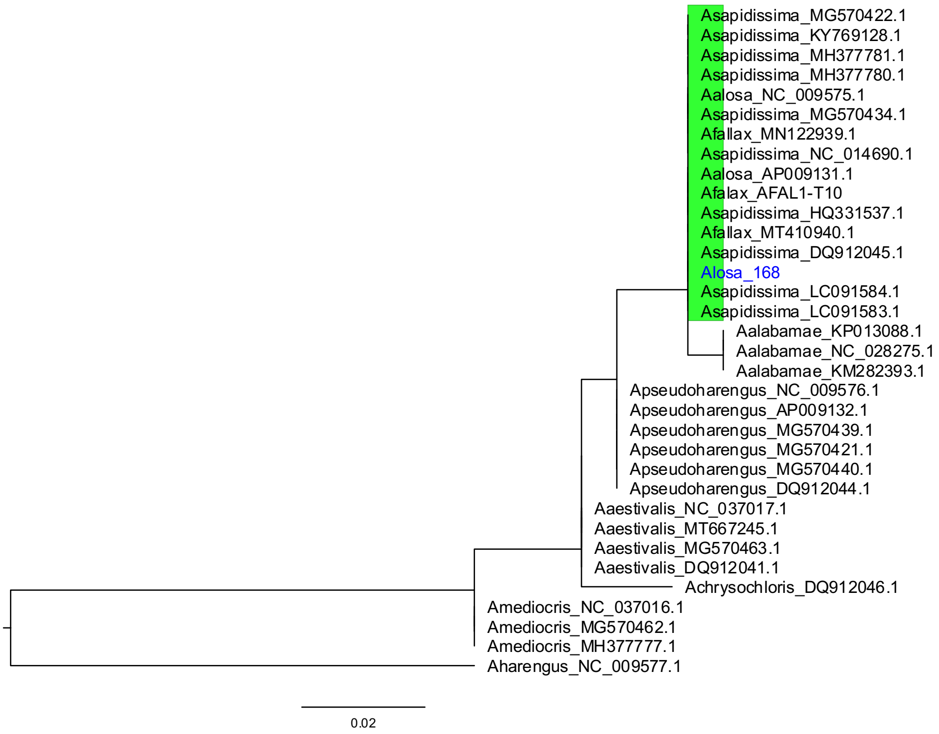


Fig. S1.1. Maximum likelihood tree of Alosa ASVs together with existing Alosa sequences in GenBank. ASV taxa labels are in blue while the clade including them is highlighted in green.

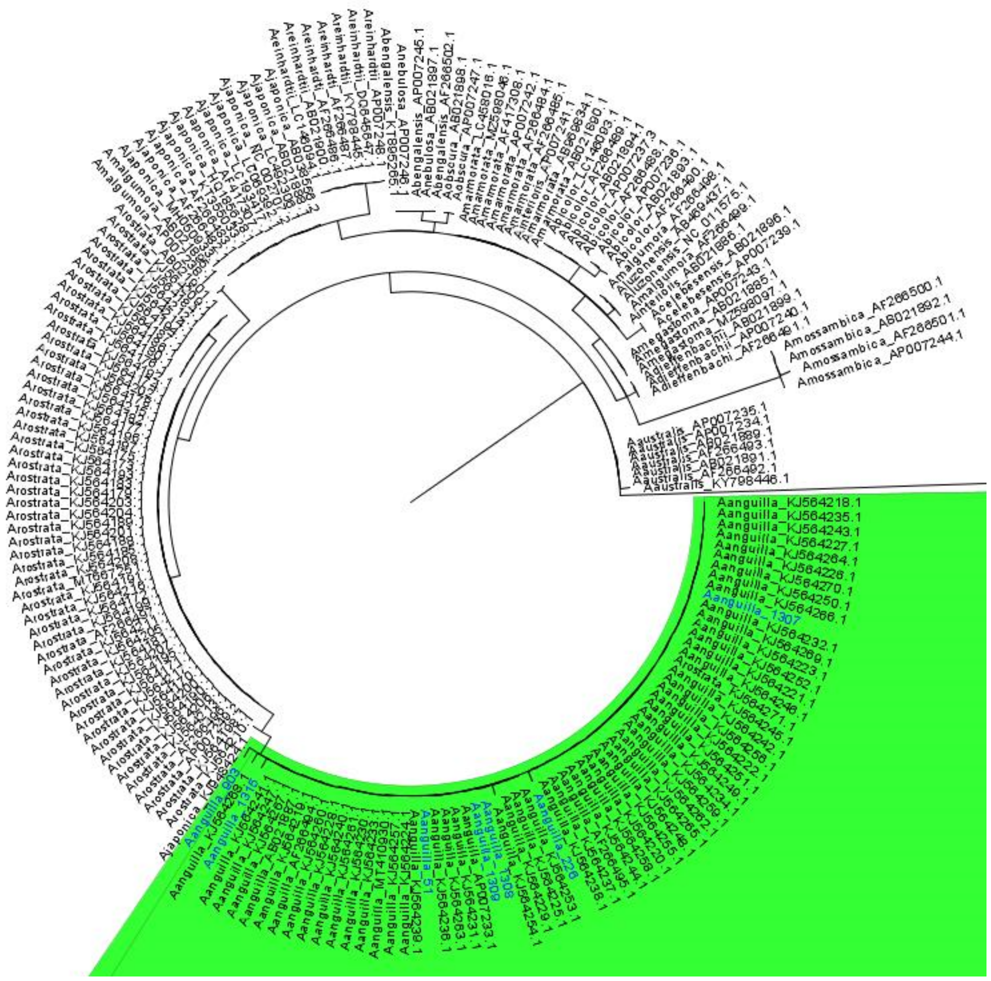

Fig. S1.2. Maximum likelihood tree of Anguilla ASVs together with existing Anguilla sequences in GenBank. ASV taxa labels are in blue while the clade including them is highlighted in green.

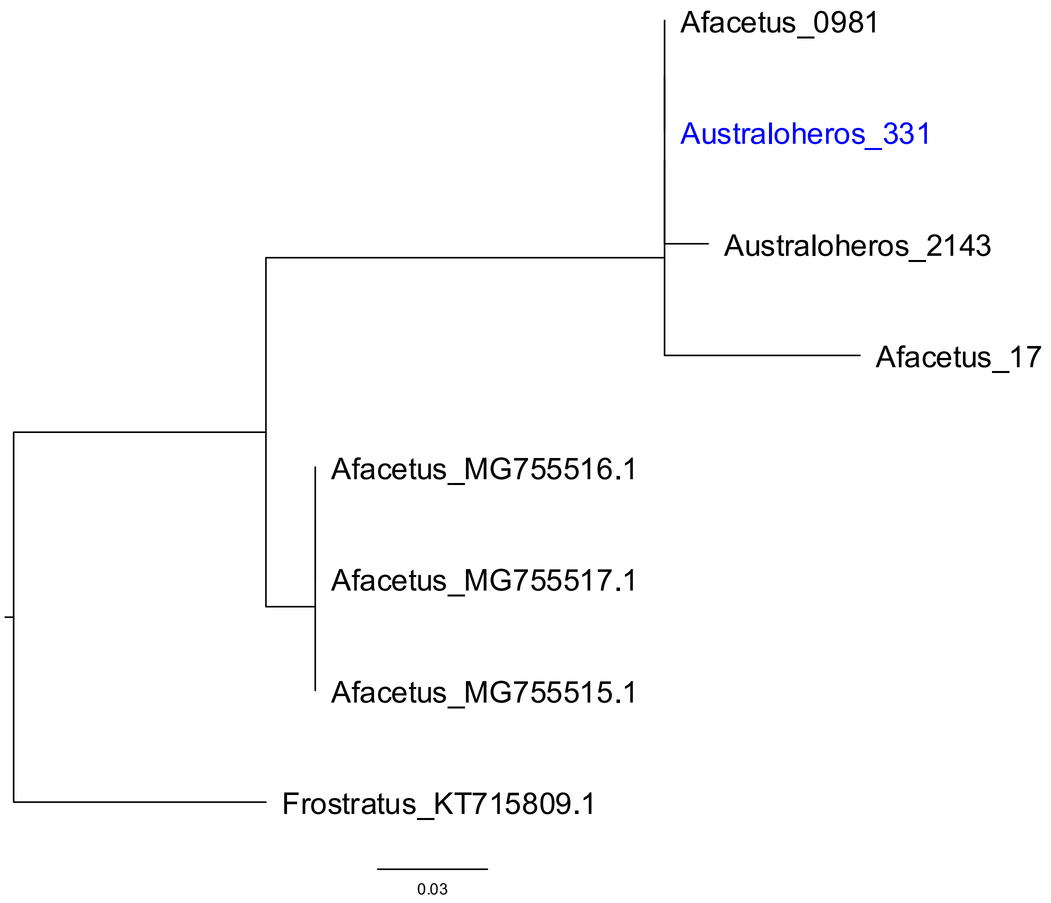


Fig. S1.3. Maximum likelihood tree of *Australoheros* ASVs together with existing *Australoheros* sequences in GenBank. ASV taxa labels are in blue.

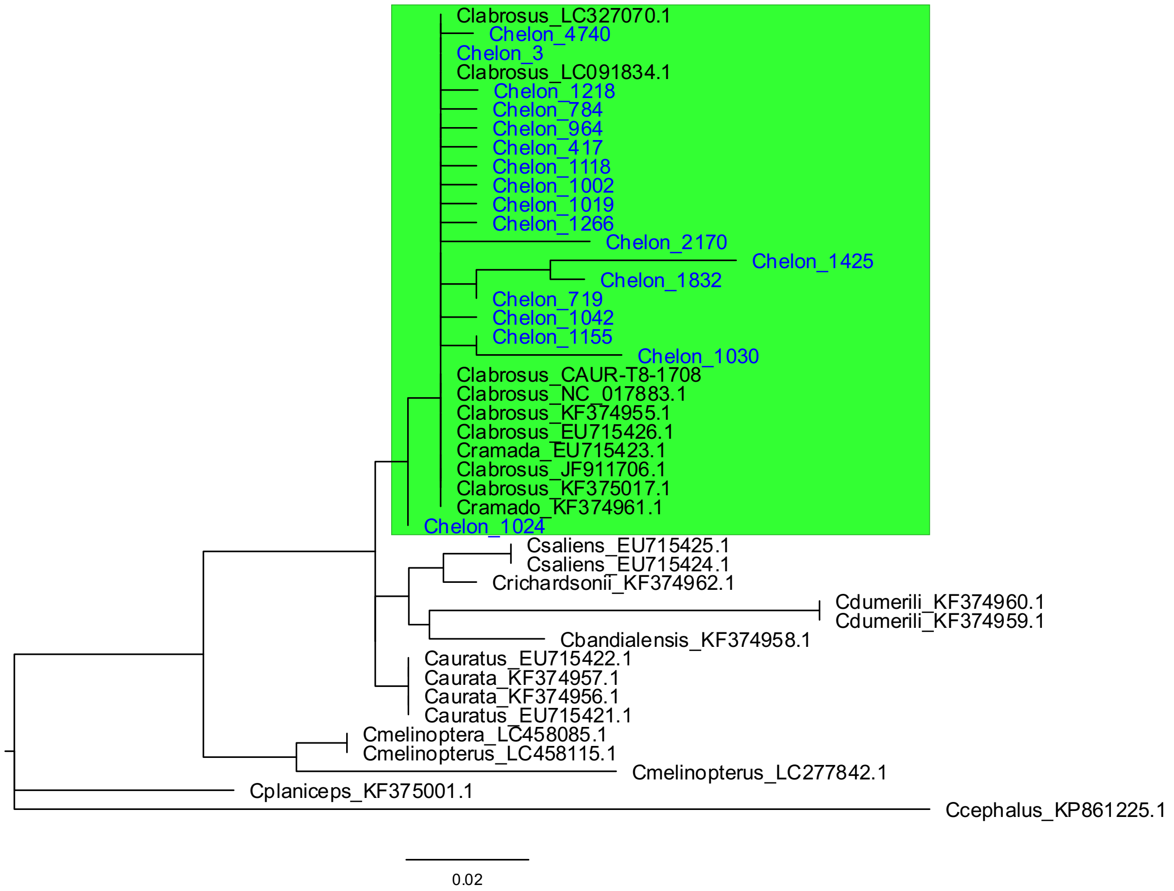


Fig. S1.4. Maximum likelihood tree of *Chelon* ASVs together with existing *Chelon* sequences in GenBank. ASV taxa labels are in blue while the clade including them is highlighted in green.

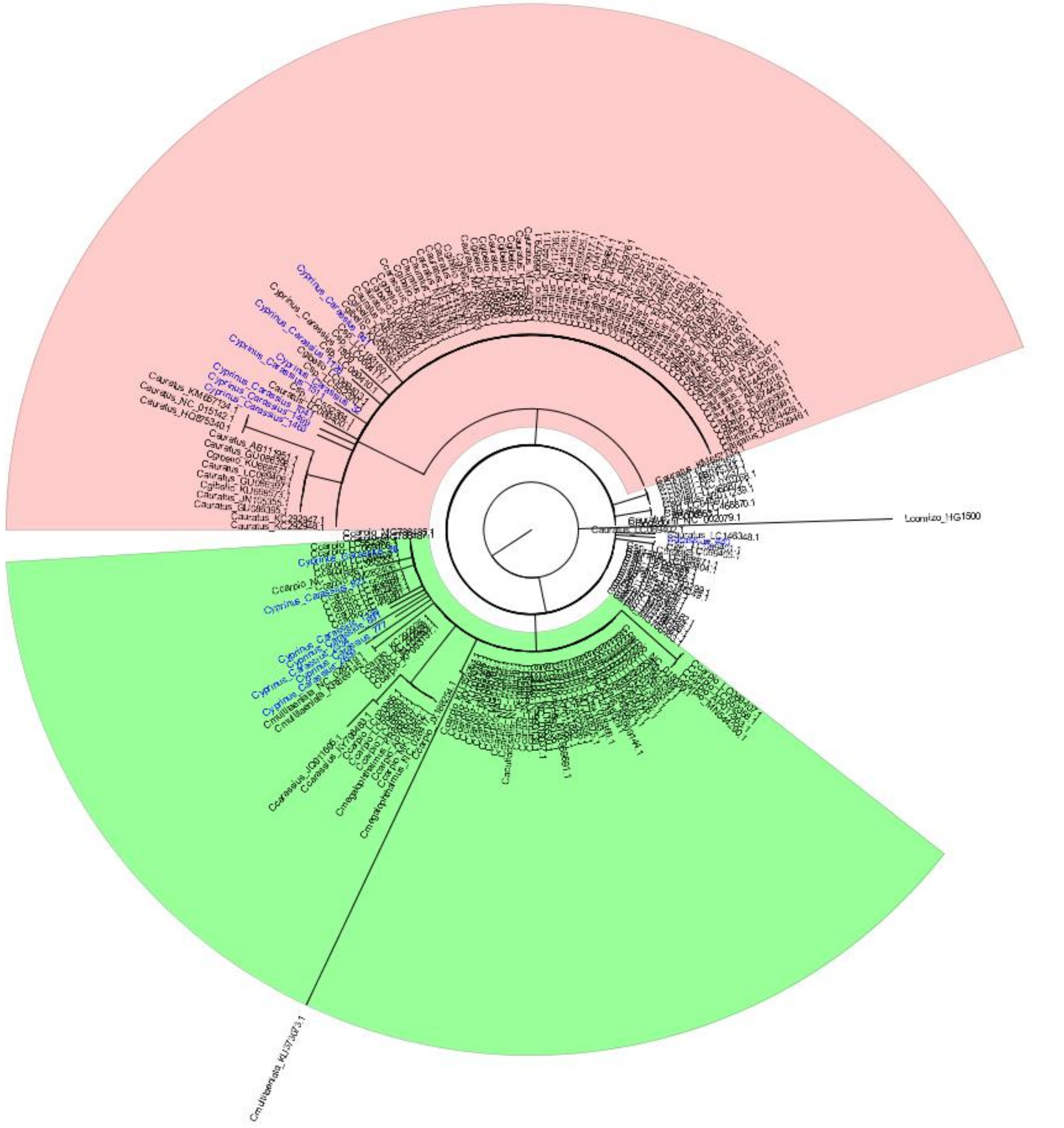

Fig. S1.5. Maximum likelihood tree of *Cyprinus* and *Carassius* ASVs together with existing sequences in GenBank. ASV taxa labels are in blue while the clade including *Carassius* ASVs is highlighted in green and *Cyprinus* in red.

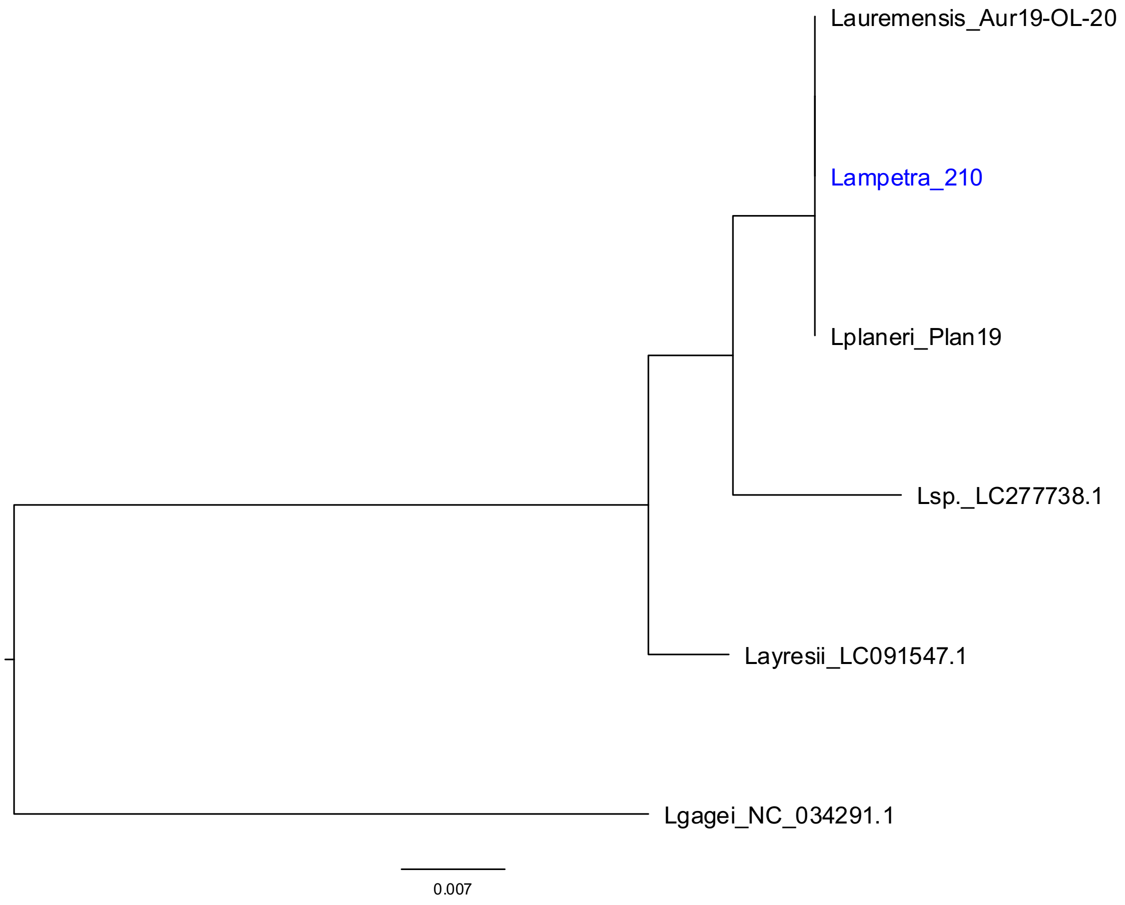


Fig. S1.6. Maximum likelihood tree of *Lampetra* ASVs together with existing *Lampetra* sequences in GenBank. ASV taxa labels are in blue while the clade including them is highlighted in green.

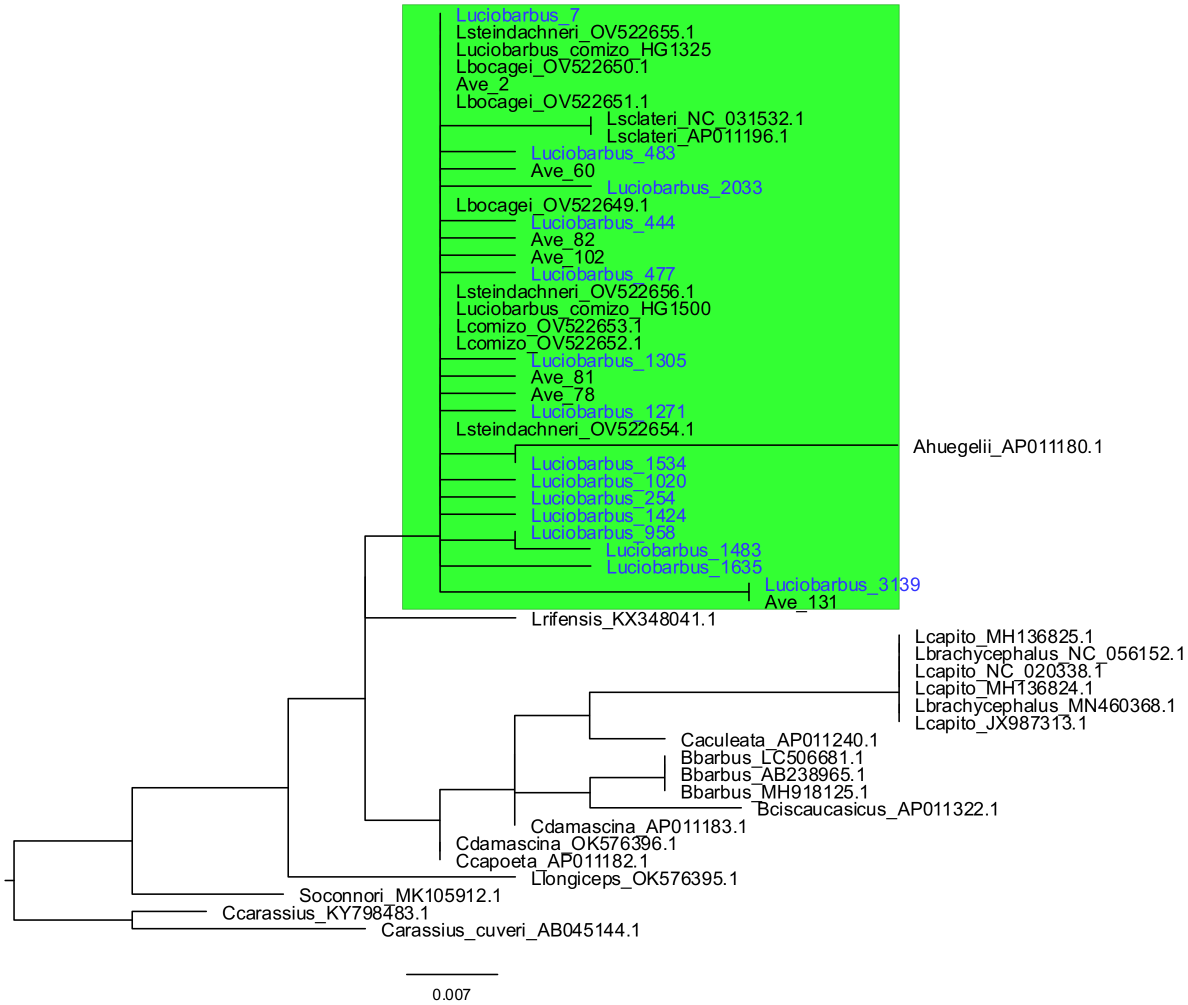


Fig. S1.7. Maximum likelihood tree of *Luciobarbus* ASVs together with existing *Luciobarbus* sequences in GenBank. ASV taxa labels are in blue while the clade including them is highlighted in green.

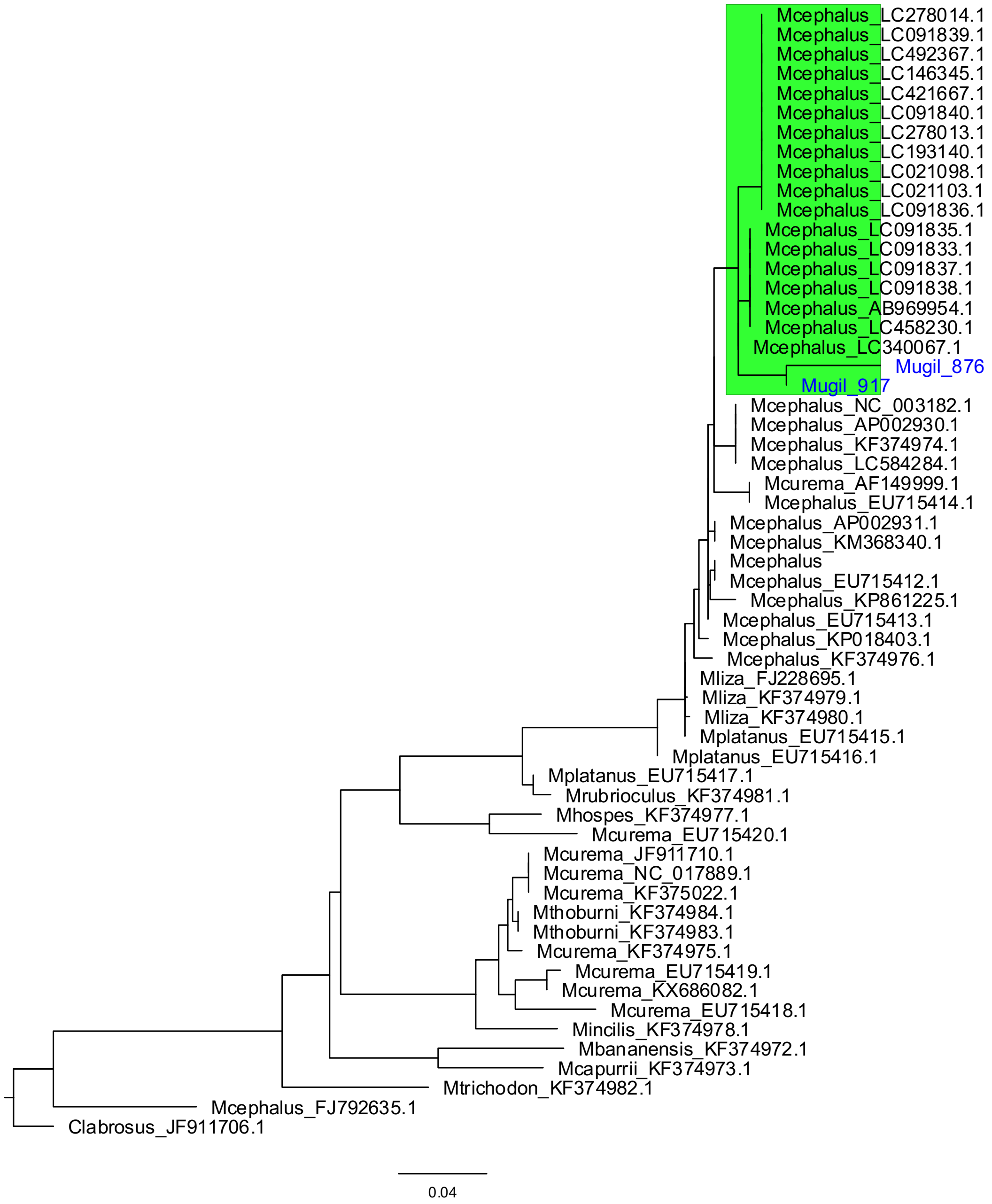


Fig. S1.8. Maximum likelihood tree of *Mugil* ASVs together with existing *Mugil* sequences in GenBank. ASV taxa labels are in blue while the clade including them is highlighted in green.

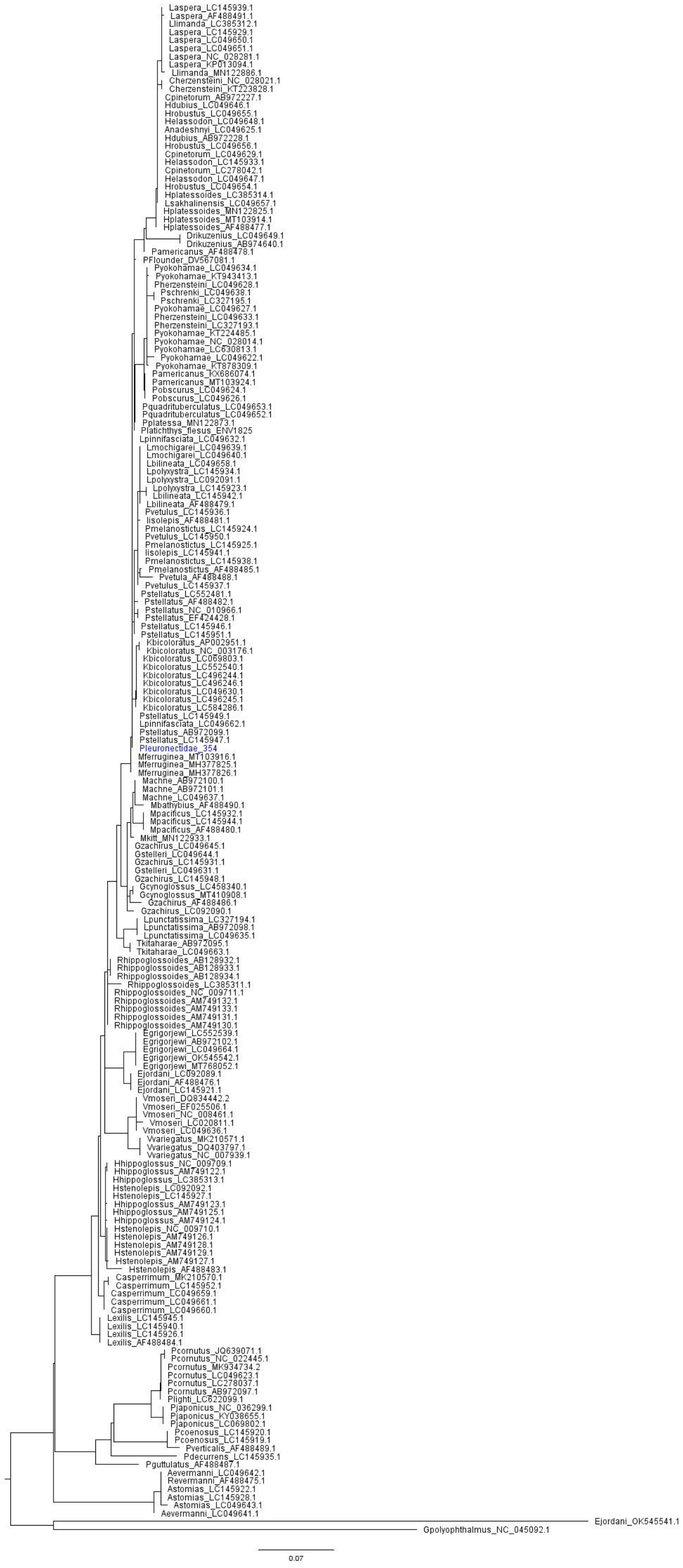

Fig. S1.9. Maximum likelihood tree of Pleuronectidae ASVs together with sequences from this family available in GenBank. ASV taxa labels are in blue.

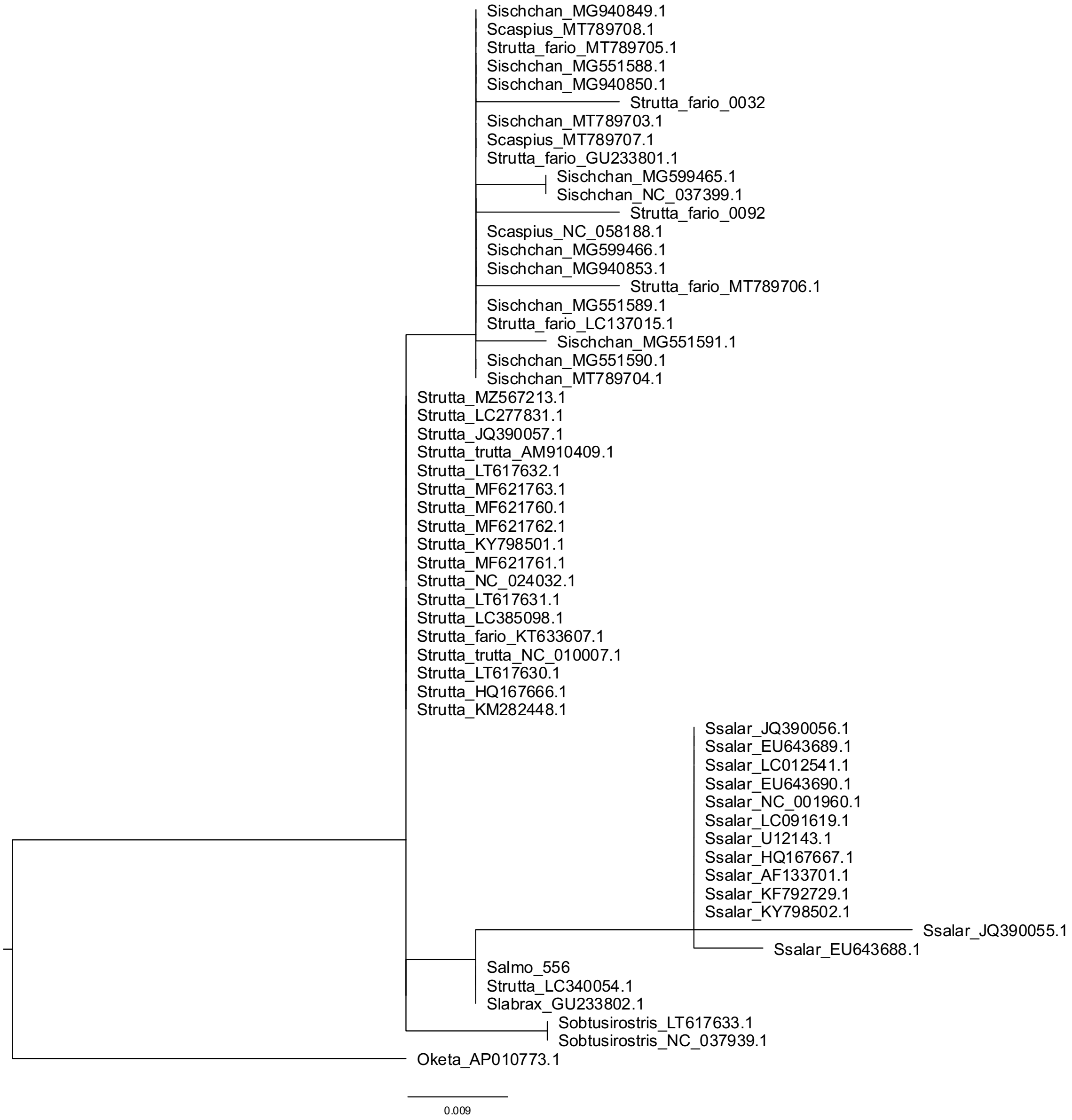


Fig. S1.10. Maximum likelihood tree of *Salmo* ASVs together with existing *Salmo* sequences in GenBank.

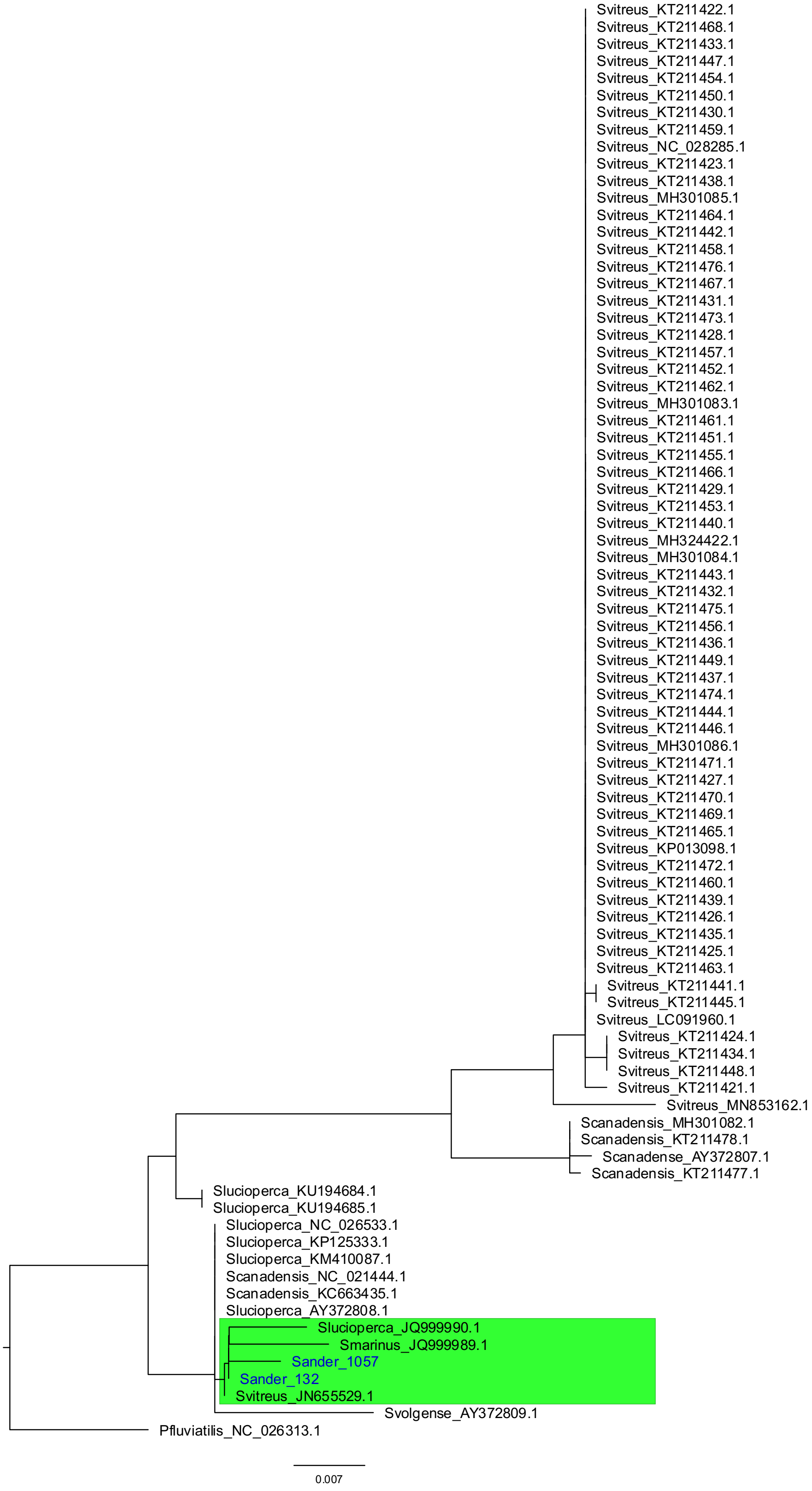


Fig. S1.11. Maximum likelihood tree of Sander ASVs together with existing *Sander* sequences in GenBank. ASV taxa labels are in blue while the clade including them is highlighted in green.

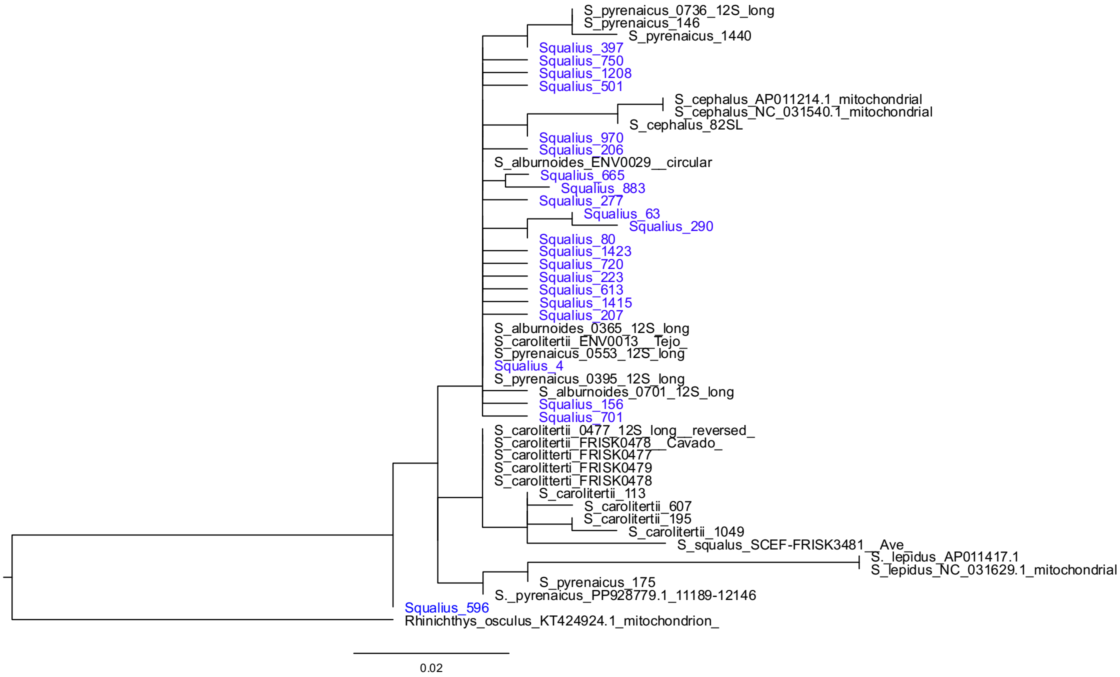


Fig. S1.12. Maximum likelihood tree of *Squalius* ASVs together with existing *Squalius* sequences in GenBank. ASV taxa labels are in blue. The phylogeny for this genus was not informative for species delimitation due to the lack of resolution and polyphyly among the included taxa.
